# Gateway: patient olfactory neurons for large-scale discovery in neurodegenerative disease

**DOI:** 10.64898/2026.06.10.731272.1

**Authors:** Kevin Zhu, Mason Sanfilippo, Minyoung Amy Oh, Mostafa Ahmed, Alex Aldrich, David Nyberg, Renan Sauteraud, Nate Dalva

## Abstract

An estimated 42% of Americans over age 55 will develop dementia, but the molecular understanding of dementia and neurodegenerative disease is constrained because the living human brain cannot be routinely sampled during disease progression. Olfactory sensory neurons provide a clinically accessible neuronal tissue source with developmental, transcriptional, and disease-relevant links to the central nervous system. Here we describe Gateway, a platform that combines device guided olfactory epithelium biopsy, onsite fixation, and 10x Genomics FLEX RNA profiling to generate single-cell transcriptomic data from living patient neurons. We present a 4-million-cell atlas representing 202 human donors, including healthy controls and individuals with neurodegenerative diseases, and release it as an open resource through CELLxGENE. We define the cellular composition of the human olfactory epithelium and show that Gateway captures neuronal functional and compartmental programs and detects more brain-enriched genes than other clinically accessible transcriptomic sample types. In exploratory analyses of Alzheimer’s Disease and Parkinson’s Disease, we identify dysregulation of pathways and GWAS-implicated genes related to key neurodegenerative mechanisms such as neuroinflammation, endolysosomal biology, proteostasis, and synaptic maintenance. Together, this atlas and clinical workflow establish living patient olfactory neurons as a scalable complementary modality for neuroscience research, target discovery, and biomarker development in neurodegenerative disease.

## Introduction

Neurodegenerative diseases (NDDs) affect more than 57 million people worldwide, and it is estimated that 42% of Americans over age 55 will go on to develop dementia (1, 2). The molecular understanding of these conditions is constrained by a simple anatomical problem: the living human brain cannot be routinely biopsied. In oncology, immunology, and many chronic diseases, disease-relevant patient tissue can be sampled directly and repeatedly. The living human brain is not so amenable. As a result, molecular studies of Alzheimer’s disease (AD), Parkinson’s disease (PD), and related disorders have depended on postmortem brain tissue, animal models, and cell models such as induced pluripotent stem cell (iPSC)-derived neurons. These models are essential, but each also has inherent limitations. Postmortem brain tissue typically reflects end-stage pathology. Animal models have induced and simplified versions of complex diseases in species that do not naturally have similar NDDs. Cell models approximate distinct components of disease pathology outside the native patient or tissue context (3, 4). With the failure rate of therapeutic programs for AD and PD among the highest in medicine, new models are warranted. A scalable source of living human neuronal tissue, which can be sampled repeatedly from the same patients, would address central limitations in NDD research and therapeutic development.

The olfactory epithelium (OE) is an exception to the inaccessibility of human neural tissue. Located in the superior nasal cavity, the OE is a neuroepithelium that contains olfactory (sensory) neurons, bipolar excitatory neurons that mediate the sense of smell and whose axons project transcranially to form synapses in the central nervous system (CNS). Olfactory neurons are the only neurons whose molecular profiles are clinically accessible without major surgery (5). Olfactory neurons are native neurons rather than surrogates or derived cells. They share developmental lineage with CNS neurons and are actively signaling excitatory neurons (6). The OE also supports continuous adult neurogenesis, with basal stem cells giving rise to intermediate neural progenitors and maturing to neurons throughout life (7–9).

The promise of human olfactory neurons has been constrained by the difficulty of accessing the OE at scale. The OE is found in a narrow region in the superior and posterior nasal cavity (10). Reaching it requires navigating variable nasal anatomy toward a small target adjacent to the delicate cribriform plate. This anatomy has historically required endoscopically guided biopsy by trained otolaryngologists, limiting the procedure, in practice, to specialized academic centers and research groups (11). As a result, olfactory neuronal tissue has remained operationally difficult to use as the basis for large, multi-site molecular studies.

The relevance of olfactory neurons to neurodegeneration is supported by several lines of evidence. Olfactory dysfunction is among the earliest symptoms of AD and PD, often preceding motor or cognitive symptoms by years. Histology of postmortem and living olfactory tissue has demonstrated that the OE accumulates hallmark neuropathology in NDD (12–18). In AD, amyloid-beta accumulations in OE have been observed to correlate with post-mortem cortical amyloid-beta burden (13). In PD, toxic alpha-synuclein has been found to be enriched compared to healthy controls by seed amplification assay and immunofluorescent imaging (17, 18). The anatomical position of the OE at the interface between the airway and the CNS, adjacent to cerebrospinal fluid (CSF) drainage pathways through the cribriform plate, further suggests that this tissue may register neuroinflammatory processes characteristic of neurodegeneration (19). A recent single-cell transcriptomic study of olfactory brush biopsies identified Alzheimer’s disease-associated immune and neuronal signatures in a preclinical and clinical AD cohort (5).

Despite these advances, previous molecular studies of human olfactory neurons have been limited in scale and clinical reach. Existing single-cell datasets from the human OE typically comprise hundreds to low thousands of neurons from fewer than 25 donors (5, 20–22). Most have also applied conventional 10x Genomics 3’ chemistry, which requires fresh tissue and immediate processing (5, 20–22). These constraints have made it difficult to assemble cohort sizes that power disease comparisons, biomarker discovery, and population-scale analyses.

Here we describe Gateway, a scalable clinical and molecular platform for generating single-cell transcriptomic data from living patient olfactory neurons, and we present the resulting atlas. The platform combines a minimally invasive nasal biopsy procedure with a novel medical device, immediate tissue fixation, and central processing with 10x Genomics FLEX fixed RNA profiling chemistry. This workflow decouples tissue acquisition from laboratory processing, enabling multi-site collections without requiring onsite single-cell infrastructure. We present an atlas of the human olfactory epithelium with 4.0 million cells spanning 202 donors, including 192 neurologically healthy controls and 10 donors with neurodegenerative disease. We define the cellular landscape of the olfactory epithelium; compare olfactory neurons to postmortem brain tissue, iPSC-derived neurons, and other accessible tissues; and provide exploratory evidence of disease-relevant transcriptomic signals. These results demonstrate a scalable platform for neuroscience research that can enable longitudinal exploration of living patient neurons.

## Results

### A minimally invasive platform for scalable olfactory neuron collection

Prior olfactory brush biopsies have been performed under endoscopic guidance by an otolaryngologist using a straight cytology brush advanced through the nasal cavity under direct visualization. This approach faces numerous practical barriers to scaling to cohort sizes needed for neurodegenerative disease research: the tortuous nasal path deflects a straight brush, the cavity constricts near the olfactory cleft, the brush is proximal to the sensitive cribriform plate, and direct visualization of the biopsy site is difficult even for experienced rhinologists.

We addressed these limitations by developing an olfactory biopsy device designed for precise, minimally invasive access to the olfactory cleft without the requirement for endoscopic visualization or subspecialist expertise. To our knowledge, this is the first specialized device for olfactory neuron collection.

Data from five separate clinical sites are included in the released atlas. Across the sites, trained otolaryngologists (one user per site) attempted olfactory biopsies with and without the Gateway device. From some donors, samples were collected from both nostrils and combined. As a result, in our overall released dataset, there are 202 total donors and 213 total samples. The atlas includes 105 donors sampled using the Gateway device and 97 donors with a standalone brush.

Sufficient neuronal yield was determined to be ≥50 mature neurons per donor. This study was not designed to benchmark the hit rate of either system. At the donor level, the neuronal hit rate of the brush was 27%. With the device, the neuronal hit rate was 71%. At the sample level, the mean and median mature neuron counts with the brush were 88 and 1, and with the device were 1,338 and 268.

### A 4-million-cell atlas of the living human olfactory epithelium

Biopsy samples were processed using the 10x Genomics FLEX assay, which uses a split probe-based chemistry to capture whole cell transcriptomes (18,129 genes) from fixed input material. Libraries were sequenced on Illumina NovaSeq X Plus instruments and aligned using Cell Ranger 10.0.0 against GRCh38 and the human 2.0 probe set reference. We processed a total of 4,028,275 cells passing quality control filters which are included in the released atlas.

The Gateway scRNAseq dataset was processed per sample through a computational pipeline invoking Cell Ranger 10.0.0 for alignment and demultiplexing; quality control filtering based on detected features, UMI counts, heterotypic doublet detection across three methods (scDblFinder, scds, DoubletFinder); and cell type annotation via label transfer from a donor diverse reference atlas built from manually curated, iteratively refined cell populations for both coarse-resolution and fine-resolution cell types (23–25). Samples were integrated across the full cohort using scANVI to generate a unified atlas to enable cross-donor, cross-batch, and disease comparisons.

We identified 11 coarse resolution cell types spanning the epithelial, immune, and neuronal compartments of the olfactory and respiratory mucosa. The atlas includes mature olfactory sensory neurons (mOSNs), immature olfactory sensory neurons (iOSNs), intermediate neuronal progenitors (INPs), sustentacular cells, microvillous cells, horizontal basal cells, Bowman’s gland cells, respiratory ciliated and secretory epithelial cells, and immune populations including T cells, macrophages, and dendritic cells. Within each major lineage we resolved sub-coarse heterogeneity by re-clustering on a 24-sample, covariate-balanced cohort using Harmony-integrated PCA and Leiden clustering, with subtype identity assigned from literature-derived marker signature scoring. Labeled cells were selected on the basis of marker fidelity, exclusivity, silhouette, and cross-validation concordance, then merged into a single integrated reference object containing 50 fine-resolution subtypes. The resulting taxonomy resolves neuronal maturation states (Early/Late INP and iOSN; Early-mature, Fully-mature, and Stressed mOSN), regenerative basal-cell states (quiescent, activated, and cycling HBC; deuterosomal ciliated precursors), tissue-immune compartments (cDC1/cDC2/pDC, classical/non-classical monocytes, tissue-resident and inflammatory macrophages, naive, effector-memory, and cytotoxic T-cell subsets), and chemosensory and ionocyte populations to support downstream quantification of disease-associated composition shifts, identification of biomarker-relevant states such as fully-mature mOSN abundance as a candidate neurodegeneration readout, and cross-study integration via the Cell Ontology. (Fig. 2)

**Fig. 1.**
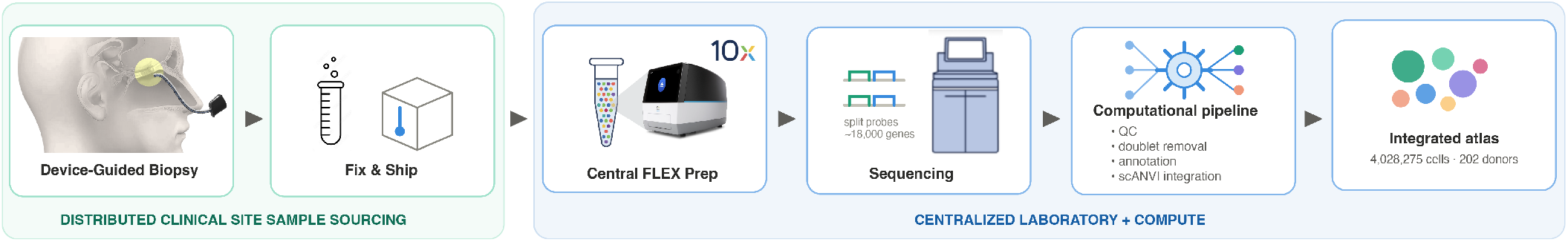
Schematic of Gateway platform and data generation process. It combines device-guided biopsy and fixation with centralized single-cell transcriptomic FLEX processing, sequencing, computational integration, and atlas generation.

**Fig. 2.**
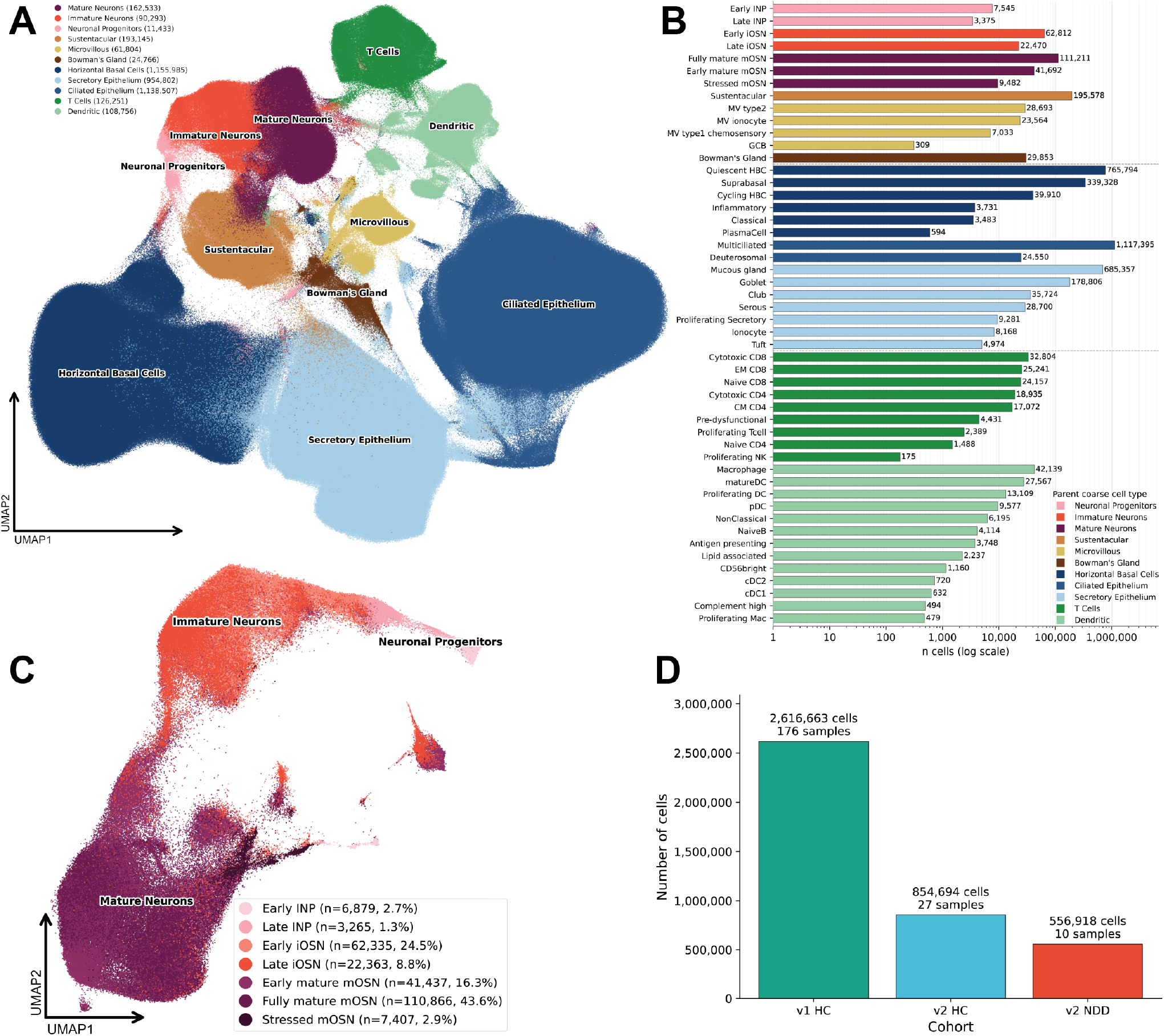
Single-cell atlas of the Gateway human olfactory epithelial cohort. (A) UMAP of 4,028,275 cells from 202 donors, colored by 11 coarse cell types spanning olfactory neuronal, basal, respiratory epithelial, glandular, immune, and other populations. (B) Fine-resolution cell-state composition across the atlas, showing 50 annotated subtypes grouped by parent coarse cell type and plotted by cell count on a log scale. (C) Cell and donor contributions by cohort, including healthy controls processed with FLEX v1 and v2 chemistry and neurodegenerative disease donors processed with FLEX v2.

In constructing the atlas, we used both 10x Genomics FLEX v1 and FLEX v2/FLEX APEX reagents as they became available and found that FLEX v2 extended the major operational advantages of the FLEX workflow while modestly improving several performance features (Supp. Table 1). FLEX’s fixed-sample, probe-based chemistry was well suited to a multi-site olfactory epithelium atlas because it enabled sample fixation at collection, centralized batch processing, reduced ambient RNA contamination, and capture of intact neuronal transcriptomes without requiring nuclear isolation. In paired donor comparisons, FLEX v2 produced highly consistent cell-type annotations, comparable library sizes and doublet rates, and slightly higher gene detection than v1. Canonical markers for major olfactory epithelium populations, including the olfactory neuron lineage, respiratory epithelial, and basal cells, remained strongly cell-type specific in v2 and in several cases showed modestly stronger enrichment. Overall, FLEX v2 provided a robust and scalable chemistry for fixed olfactory epithelial single-cell profiling, supporting reliable recovery of diverse epithelial, immune, stromal, and neuronal-lineage populations across the atlas.

**Table 1.** Gateway cohort demographics and sample composition.

| Characteristics | Overall | Healthy | Neurodegenerative |
| --- | --- | --- | --- |
| Donors, n | 202 | 192 | 10 |
| Samples, n | 213 | 203 | 10 |
| Cells, n | 4,028,275 | 3,471,357 | 556,918 |
| Age, median (IQR) | 64 (47–72) | 64 (46–72) | 74 (62–79) |
| Age not reported | 5 | 5 | 0 |
| Sex, n (%) |  |  |  |
| Male | 82 (40.6%) | 77 (40.1%) | 5 (50%) |
| Female | 118 (58.4%) | 113 (58.9%) | 5 (50%) |
| Not reported | 2 (1.0%) | 2 (1.0%) | 0 |
| Race and Ethnicity, n (%) |  |  |  |
| White | 67 (33.2%) | 61 (31.8%) | 6 (60%) |
| Asian | 56 (27.7%) | 56 (29.2%) | 0 |
| Hispanic/Latino | 20 (9.9%) | 20 (10.4%) | 0 |
| African American | 18 (8.9%) | 17 (8.9%) | 1 (10%) |
| Multiracial | 20 (9.9%) | 18 (9.4%) | 2 (20%) |
| Not reported | 21 (10.4%) | 20 (10.4%) | 1 (10%) |
| Collection method, donor n (%) |  |  |  |
| Brush | 97 (48.0%) | 97 (50.5%) | 0 |
| Device | 105 (52.0%) | 95 (49.5%) | 10 (100%) |

### Olfactory neurons are accessible living neuronal tissue

To validate living olfactory neurons for neuronal research, we benchmarked our data against multiple reference datasets including direct CNS tissue and accessible peripheral surrogates. We compared Gateway mOSNs to excitatory neurons from the postmortem and living brain (SEA-AD DLPFC and MTG, Vornholt et al. Living Brain Project) (26), iPSC-derived excitatory neurons (Matassa et al.) (27), mOSNs from published olfactory epithelium datasets (Durante et al., Olivia et al., D’Anniballe et al.) (5, 21, 22), fibroblasts (Tabula Sapiens) (28), and biofluid transcriptomics (Schafflick et al.) (29). All datasets were processed uniformly, subsetted to relevant cell populations (healthy donors, excitatory neuron subtypes when neuronal), and converted to common gene identifiers to maximize comparability.

We defined a brain-enriched gene set as the 1,178 genes whose normalized expression in the brain exceeds their expression in any other tissue by at least four-fold, using GTEx Consortium tissue-level expression data (30). We then quantified detection of these brain-enriched genes at progressively stringent per-cell prevalence thresholds. Living brain single-cell transcriptomic data detected 632 brain-enriched genes in 10% of excitatory neurons. Postmortem brain datasets set the upper bound and detected 886 genes in at least 10% of excitatory neurons.

Gateway olfactory neurons provided greater brain-gene visibility than other accessible sample types which may otherwise be used for longitudinal human studies. Gateway mature neurons with FLEX v2 detected 974 brain-enriched genes overall and retained 619, 540, and 333 brain-enriched genes at the 5%, 10%, and 30% of cells, respectively. CSF and blood-derived cells tested had fewer than 50 genes detected at the 10% of cells threshold. iPSC-derived neurons, which require ex vivo re-engineering, also had slightly lower brain-gene visibility than olfactory neurons with 510 genes at the 10% of cells threshold. (Fig. 3) Gateway samples with FLEX v1 chemistry, which were on average sequenced less deeply than with v2, detected 835 brain-enriched genes overall and retained 473, 356, and 146 at the 5%, 10%, and 30% thresholds, respectively.

**Fig. 3.**
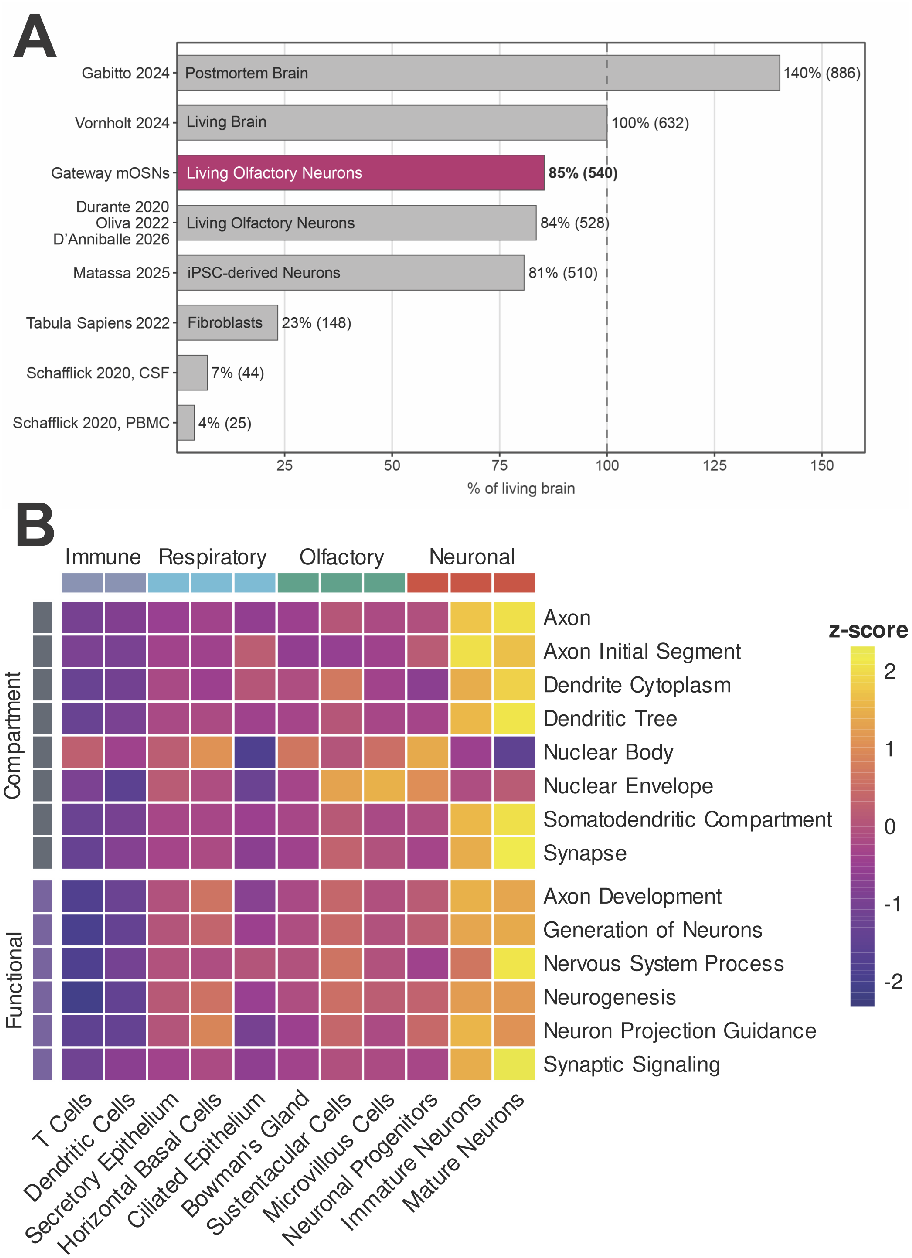
Olfactory neurons are a strong clinically accessible CNS surrogate by gene visibility. (A) Brain-enriched gene detection (1,178 genes) in at least 10% of cells by data source: Gateway mature neurons sequenced with FLEX v2 detect a median 974 brain-enriched genes/sample, reaching 85% of living-cortex detectability at the 10%-of-cells threshold, slightly higher than iPSC-derived neurons and meaningfully higher than fibroblasts and biofluids. (B) Within-atlas validity of the curated neuronal compartment/functional panel: olfactory-neuronal cell types score highest on axon/synapse/dendrite axes while non-neuronal cell types invert as expected.

### Gateway includes neuronal functional and compartmental programs

Detection of individual brain-enriched genes does not alone establish that olfactory neurons express these genes in functionally coherent patterns. To assess functional biology, we scored cells using Gene Ontology Biological Process (GOBP) modules corresponding to core neuronal programs: axon biology, synaptic function, neuronal signaling, and neuronal projection (31, 32). Gene set scores were computed at the single-cell level using a rank-based normalized Mann-Whitney U statistic (UCell), yielding a library-size-independent measure that is comparable across datasets and samples (Methods).

Within the Gateway data, immature and mature olfactory sensory neurons were enriched across neuronal modules and showed expected maturation-associated differences. Mature neurons scored higher than immature neurons for Synaptic Signaling (mature 0.083, immature 0.073, paired Wilcoxon *p <* 0.00001; *n* = 159 healthy donor samples) and Nervous System Process modules (mature 0.064, immature 0.053, *p <* 0.00001), supporting a stage-dependent increase in neuronal functional specialization across the olfactory lineage (Fig. 3).

We also scored cells using Gene Ontology Cellular Component (GOCC) modules corresponding to neuronal subcellular structures: dendrite, soma, nucleus, axon, and synapse (31, 32). The results confirmed the functional biology findings: olfactory neurons show approximately 3-fold stronger neuronal compartment enrichment than fibroblasts while preserving nuclear gene coverage comparable to postmortem cortical neurons.(Supp. Fig. 1) Within Gateway data between cell types, neurons (mature and immature) ranked most highly for every neuron-specific GOCC compartment module axon (mOSN z = +2.07; iOSN z = +1.76), axon initial segment (+1.72; +2.05), dendrite cytoplasm (+1.89; +1.48), dendritic tree (+2.12; +1.58), somatodendritic compartment (+2.08; +1.62), and synapse (+2.14; +1.46) and were de-enriched for the nuclear body module (mOSN z = −1.5; iOSN z = −0.43). (Fig. 3)

Across curated neuronal functional and compartmental modules, brain-derived neuronal datasets generally scored highest, with postmortem cortex showing the strongest overall module scores. Gateway v2 healthy donor mOSNs occupied a neuronal range similar to in-vitro engineered iPSC-derived neurons. Across GOBP functional modules, Gateway mOSNs reached a median of approximately 92% of the iPSC-derived neuron score, ranging from 72–132% across individual modules. Relative performance was strongest for Nervous System Process (132% of iPSC-derived neuron score) and Synaptic Signaling (118%). Across GOCC compartment modules, Gateway v2HC mOSNs reached a median of approximately 91% of the iPSC-derived neuron score, ranging from 54– 105%. Relative to fibroblasts, Gateway v2HC mOSNs showed enrichment across neuronal functional and compartmental modules except only nuclear body compartment (median 1.6-fold). These results indicate that Gateway mOSNs have similar functional and compartmental module neuronal signal to widely applied in-vitro neuronal models and are enriched over fibroblasts as non-neuronal substitutes.

### Neurodegenerative disease programs in Gateway

We next asked whether living olfactory neurons capture molecular programs relevant to neurodegenerative disease (NDD). The disease cohort includes 5 donors with clinically diagnosed Alzheimer’s disease and 5 donors with clinically diagnosed Parkinson’s disease. All neurodegenerative disease (NDD) patients were sampled with the Gateway device and FLEX v2 chemistry, as were the healthy control donors used in this analysis.

We examined whether Gateway olfactory epithelial cell types express genes implicated in major NDD pathways across all donors. We declared *a priori* a list of 23 canonical NDD genes, considering Alzheimer’s disease, Parkinson’s disease, frontotemporal dementia, and amyotrophic lateral sclerosis (ALS). (Fig. 4) Across neuronal and non-neuronal cell types, we detected broad expression of genes associated with synuclein biology, amyloid processing, tau biology, lysosomal function, mitophagy, ubiquitin-proteasome biology, and lipid-immune risk. Several canonical NDD genes, including SNCA, APP, BIN1, PINK1, and UCHL1, were detected at the highest mean expression in neuronal cell types. (Fig. 4)

**Fig. 4.**
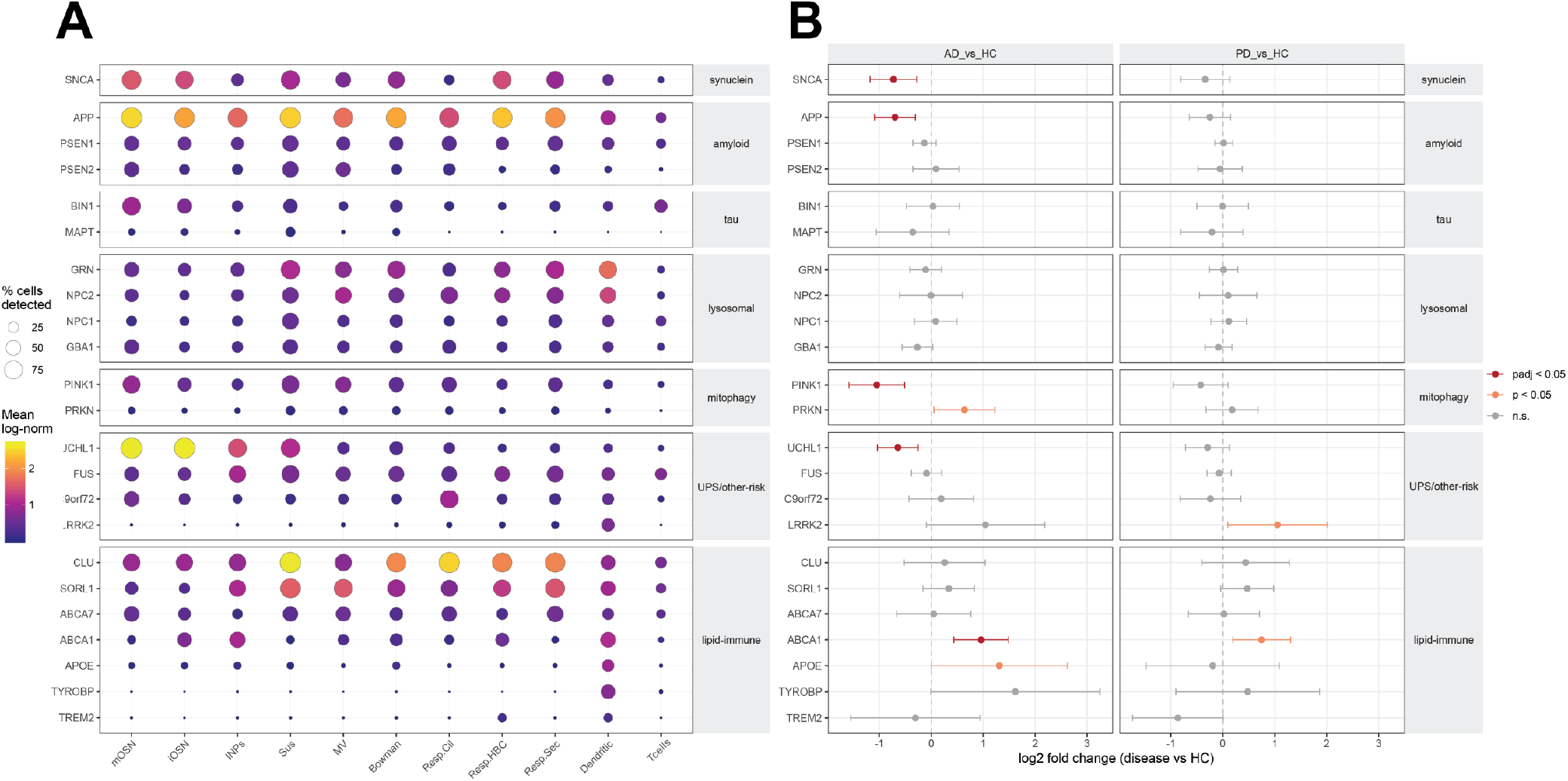
Expression and disease-associated perturbation of canonical neurodegenerative disease genes in Gateway olfactory epithelial cell types in FLEX v2 data. (A) Dot plot showing detection frequency and mean expression of selected genes across Gateway cell types. (B) Differential expression of the same gene panel in mature olfactory neurons from Alzheimer’s disease (n=5) and Parkinson’s disease (n=5) donors compared with healthy controls (n=27).

We computed differential expression (DESeq2) on cell type level pseudobulks with covariates for donor age, sex, and library size against 27 cognitively normal donors. All Gateway data in this analysis were produced with FLEX v2 chemistry. We compared expression of the 23 *a priori* selected genes in fully mature olfactory neurons of each disease versus cognitively normal controls. The analysis included 22,943 NDD fully mature neurons (8,136 AD, 14,807 PD) and 44,875 control fully mature neurons. In Alzheimer’s disease, Gateway mature neurons had differential expression of SNCA (−0.73, padj = 0.007), APP (−0.7, padj = 0.007), PINK1 (−1.05, padj = 0.003), UCHL1 (−0.64, padj = 0.007), and ABCA1 (0.96, padj = 0.004) with multiple testing correction for the 23-gene panel. In Alzheimer’s disease, PRKN (0.64, p = 0.033, padj = 0.13) and APOE (1.31, p = 0.049, padj = 0.15) reached unadjusted significance at p < 0.05. In Parkinson’s disease, ABCA1 (0.75, p = 0.009, padj = 0.2) and LRRK2 (1.05, p = 0.031, padj = 0.36) reached nominal significance but were not significant after multiple testing correction. (Fig. 4)

We ran whole transcriptome differential expression in fully mature Gateway neurons, comparing the 10 NDD donors (AD and PD combined) to cognitively normal donors (n=27) to improve statistical power and examine shared neurodegenerative mechanisms. Gene set enrichment analysis (GSEA) identified a coherent upregulation in neuroinflammatory pathways. Multiple disease relevant pathways were transcriptionally downregulated in GSEA, including ubiquitin-proteasome, mitophagy, endolysosomal, amyloid fibril, and synaptic pathways. Thus, the dominant disease-associated signature in this pilot cohort is a broad shift toward inflammatory activation with concomitant reduction of protein homeostasis and synaptic maintenance programs. (Fig. 5)

**Fig. 5.**
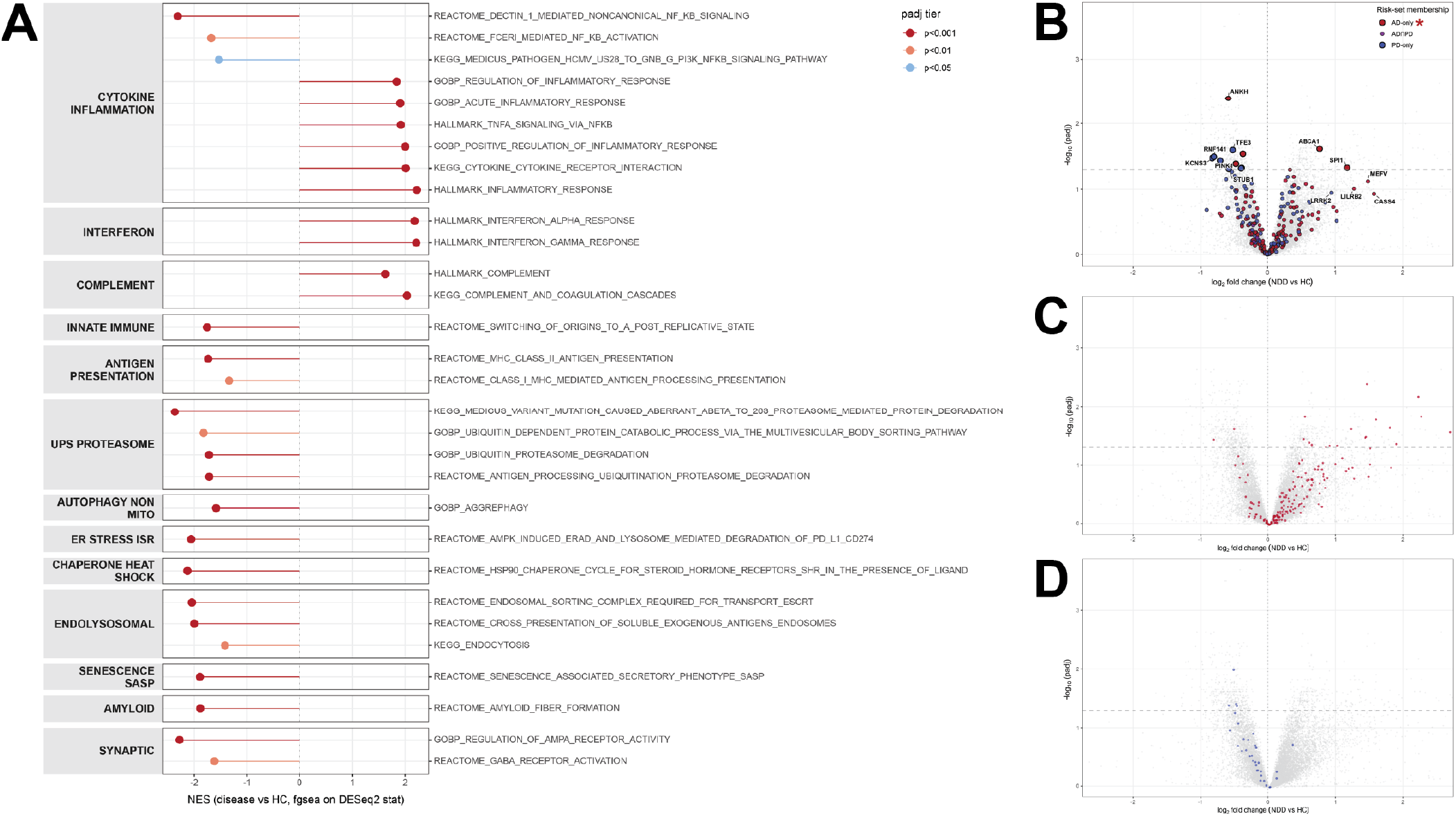
Neurodegenerative disease-associated pathway and genetic-risk signatures in Gateway neurons. (A) Gene set enrichment analysis of mature olfactory sensory neurons from neurodegenerative disease donors versus healthy controls identifies upregulation of inflammatory, interferon, and complement pathways and downregulation of proteostatic, endolysosomal, amyloid, and synaptic programs. (B) Consensus AD and PD GWAS hits overlaid on the NDD differential expression results; Alzheimer’s disease GWAS genes are significantly enriched among disease-associated transcriptional changes (NES = 1.236, p = 0.036). (C,D) Leading upregulated and downregulated pathways from panel A, respectively interferon gamma response (C) and mutation-aberrant Aβ to 26S proteasome-mediated protein degradation (D).

Finally, we asked whether genes nominated by human GWAS of AD and PD were non-randomly represented among diseaseassociated transcriptional changes in Gateway neurons. By GSEA, risk allele-associated genes (by original source mapping) for both Alzheimer’s and Parkinson’s disease were enriched in the Gateway mature neuron NDD differential expression analysis. AD-associated GWAS genes were enriched (p = 0.036) (33), with several risk genes significantly differentially expressed at the whole transcriptome level: ABCA1 (log2FC = +0.772, padj = 0.004), SPI1 (+1.177, 0.047), ANKH (−0.585, 0.004), SORT1 (−0.370, 0.029), APP (−0.474, 0.041), and ILRUN (−0.390, 0.047). Likewise, PD-associated genes were enriched (p = 0.009) (34), with significantly differentially expressed genes: RNF141 (log2FC =-0.796, padj = 0.032), KCNS3 (−0.822, 0.033), PINK1 (−0.704, 0.036), LRP10 (−0.397, 0.048). (Fig. 5)

These results provide evidence that Gateway data from patient olfactory neurons detect disease-relevant molecular programs in NDD. While exploratory, this analysis establishes that neurodegenerative disease risk genes are visible in the sampled tissue and that a small combined AD/PD cohort shows coherent alterations in inflammatory, proteostatic, endolysosomal, and synaptic pathways.

## Discussion

The central limitation in translational neurodegeneration research is that the disease tissue is unavailable while disease is unfolding. Here we present Gateway, a platform for generating molecular data from patient olfactory neurons, and release 4.0 million single cells sequenced from 202 donors. Olfactory neurons are native neurons, continuously regenerate in adulthood, and can be accessed through the nasal cavity with the Gateway medical device. The study establishes that olfactory neurons can be collected, fixed at the point of care, centrally processed, and analyzed at a scale that has not previously been practical for human neuronal tissue. This platform provides a window into patient neuronal biology.

The atlas resolves the major epithelial, immune, basal, glandular, and neuronal populations of the human olfactory mucosa, including maturation states across the olfactory neuronal lineage. We show that mature olfactory neurons express approximately 84% of brain-enriched genes compared to brain neurons from living donors, and substantially exceed alternative clinical tissue sources such as fibroblasts (23%) and CSF- and blood-derived cells (4-7%). Olfactory neurons showed coherent enrichment for neuronal functional and subcellular programs, including synaptic, axonal, dendritic, and neuronal signaling modules. Synaptic functional and compartmental modules were particularly strongly expressed in Gateway olfactory neurons, more so than in iPSC-derived neurons, potentially as a result of the whole-cell sequencing enabled by the platform instead of single-nucleus sequencing.

The neurodegenerative disease analyses provide a test of whether this modality can detect neurodegeneration-relevant biology. Olfactory neuron single-cell sequencing is an increasingly important research tool for Alzheimer’s disease (AD) and Parkinson’s disease (PD) (5). Mature neurons expressed canonical neurodegenerative disease-related genes found in CNS neurons such as SNCA (encoding alpha-synuclein), APP (encoding amyloid precursor protein), BIN1, PINK1, and others. In an exploratory cohort of donors with AD and PD compared to cognitively normal donors, with adjustments for age and sex, mature olfactory neurons were dysregulated along innate inflammatory (up), endolysosomal, amyloid, mitophagy, and synaptic pathways (down). Genes associated with AD and PD GWAS-nominated risk alleles were also enriched among disease-associated transcriptional changes.

The Gateway platform and data offer a new tool for neuroscience drug discovery and development. Neurodegenerative disease therapeutic programs often pursue targets based on genetics, postmortem tissue, animal models, and/or iPSC-derived systems. These sources leave uncertainty about whether implicated pathways are active in living patients at clinically relevant stages. First, in complex, multifactorial diseases like NDDs, longitudinal data from living patients can provide additional evidence in support of purported causative pathologies. Second, olfactory neuron data may provide additional biomarkers for patient classification or clinical trial outcomes. The Gateway workflow is compatible with many observational and interventional clinical studies. These translational tools may be most useful for therapeutic programs involving proteostasis, lysosomal and endosomal biology, innate immune signaling, synaptic maintenance, mitochondrial stress, and genetically nominated risk pathways.

The study also has broader implications for neuroscience. The human olfactory epithelium is a dynamic neural tissue containing basal stem cells, neuronal progenitors, immature neurons, mature neurons, support cells, immune cells, and glandular populations. Its accessibility creates opportunities to study adult neurogenesis, neuronal maturation, epithelial-immune interactions, environmental exposure, aging, and disease-associated neuronal and glial stress in humans. We are hopeful that this atlas, freely available in its entirety, can accelerate some of these efforts.

Several limitations should guide interpretation, dataset use, and future work. The biopsy device and workflow were developed to enable scalable collection, but this study was not designed as a formal device-performance or safety trial. The atlas includes samples generated across chemistries, sites, collection methods, and donor groups, and although computational integration and focused FLEX v2 analyses mitigate these variables, residual technical and site effects remain possible; these variables are provided in the released data. As with all RNA sequencing analysis, any interpretation should consider that the data measures transcript abundance, not proteins or spatial organization. Finally, the NDD donor population here is intentionally exploratory in size.

In summary, Gateway establishes a scalable approach for generating single-cell molecular data from living human olfactory neurons and provides a large public atlas of the human olfactory epithelium. The results support olfactory neurons as an accessible and disease-relevant complement to CNS tissue and other disease models. The exploratory neurodegenerative disease findings justify larger, clinically powered studies to test whether this tissue can support target validation, biomarker discovery, patient stratification, and pharmacodynamic monitoring. By making living patient neurons more accessible to systematic study, Gateway offers a practical new substrate for translational neuroscience.

## Methods

### Study design, participant enrollment, and sample collection

Donors were enrolled at five clinical sites over 10 months under IRB-approved protocols (Sterling IRB-13038, Sterling IRB-14403) and provided informed consent. Participants were recruited via direct clinic referral, collaborating neurologists or memory clinics, or a Sponsor-contracted recruitment partner, and enrolled after eligibility confirmation against IRB-approved materials. Eligible donors had fluency for self-report assessments and no bilateral nasal abnormalities precluding access or allergy to the topical decongestant or anesthetic.

Oxymetazoline hydrochloride 0.05% and tetracaine 2% were administered intranasally prior to collection. No serious adverse events were reported. Observed events, including minor epistaxis, local discomfort, sneezing or lacrimation, and vasovagal response, were self-limited.

### 10x Genomics FLEX library preparation

All samples were fixed in the clinic and shipped to a central processing laboratory. Fixed olfactory epithelial samples were processed using the 10x Genomics Chromium Flex workflow according to the manufacturer’s protocols: Flex v1, CG000787; Flex v2, CG000834. Target input was 20,000 cells per sample for GEM generation on a Chromium X.

### Sequencing and count matrix generation

FLEX libraries were sequenced on an Illumina NovaSeq X Plus instrument. Library depth was targeted for a minimum of 10,000 reads per cell, with achieved sequencing depth varying by cohort and sequencing run.

Raw sequencing data were processed using Cell Ranger 10.0.0, using the included GRCh38 reference and Human Transcriptome v2.0.0 GRCh38 2024 A probe set. The `multi` command was used for probe-barcode demultiplexing and per-sample count matrix generation, while the `aggr` command was used to combine sample-level outputs across GEM wells where applicable.

### Sample processing and annotation

Per-sample FLEX outputs were processed using a custom scRNA workflow based largely on Seurat. Cell Ranger filtered count matrices were loaded into Seurat objects, Hao et al., 2023, for standardized processing including SCTransform, PCA, UMAP, nearest-neighbor graph construction, and Leiden clustering (35).

As human olfactory epithelium has not been sequenced using 10x FLEX methodology, we constructed a custom reference for cell-type annotation from samples present in the released atlas. Initial clustering of the samples revealed clusters whose cell-type proportions and transcriptional markers aligned with published human olfactory epithelium data (22). We constructed a 24-donor cohort built to represent donor diversity across age, sex, race, and disease conditions.

Cells from this cohort underwent normalization to 10,000 counts per cell, log-transformation, selection of 3,000 highly variable genes, scaling, PCA projection, Harmony integration by donor key, UMAP embedding, and Leiden clustering to generate a reference. From this reference cohort, we subset the top 25% of cells from these coarse-level populations based on the number of UMIs per cell, detected genes per cell, and singlet status in order to generate a reference cell population for subsequent coarse cell-type annotation. Subclustering of coarse populations and marker calling was used to derive the 50 fine-level cell types. The resulting fine labels were transferred at the single-sample level using expression-based k-nearest-neighbors classification.

### Atlas construction

The released atlas contained 4,028,275 cells from 202 donors represented by 213 final sample identifiers. Cells were retained after cohort-level QC requiring adequate UMI depth, detected genes, mitochondrial fraction, doublet status, and coarse cell-type prediction confidence. Specifically, retained cells satisfied `nCount_RNA` ≥ 500, `nFeature_RNA` ≥ 250, `percent.mito` *<* 20, were not determined to be doublets by a majority of the doublet-calling programs, and had coarse cell-type prediction confidence *≥* 0.6 from Seurat’s `TransferData` function.

The final expression matrix profiles cells across an 18,127-gene feature space. PAXX, ENSG00000148362, and SWIM2, ENSG00000139656, were removed from the FLEX panel to match the data made available on CELLxGENE(36, 37). Metadata retained for downstream analyses included donor identifier, sample identifier, sex, age, race/ethnicity, disease condition, FLEX chemistry version, coarse cell-type labels, fine cell-type labels, device usage, and QC metrics.

A low-dimensional representation was constructed for visualization using scVI and scANVI. The initial scVI embedding was constructed with donor as a batch key in a 50-dimensional latent space. scANVI was then initialized from the trained scVI model and fine-tuned using the coarse cell-type labels. The trained model was used to infer the 50-dimensional latent representation for all cells in the final atlas. A k-nearest-neighbors graph with non-default parameter `n_neighbors = 100` was constructed from the scANVI latent representation using cosine distance and used for generation of a UMAP visualization with non-default parameters `min_dist = 0.8` and `spread = 1.8`.

### External dataset curation

External datasets were curated to benchmark Gateway olfactory neurons against direct CNS tissue and other clinically accessible sample types. Comparator datasets included published human olfactory epithelial datasets from the Goldstein laboratory; SEA-AD postmortem brain; living brain gene expression from the Living Brain Project, Vornholt et al., 2023; Tabula Sapiens fibroblasts; Matassa et al. iPSC-derived neural organoids; and Schafflick et al. biofluid single-cell datasets from CSF and PBMC (5, 21, 22, 26, 27, 29, 38).

Datasets were loaded into Seurat v5 objects, restricted to healthy donors, and subset to the relevant comparable cell types: excitatory neurons for neuronal datasets and all cells for fibroblasts, CSF, and PBMC. Datasets were harmonized to Ensembl gene identifiers. Gene detection at the cell level was defined as a count value greater than 0.

### Brain-enriched gene detection benchmark

Brainenriched genes were defined using the Human Protein Atlas-derived RNA expression consensus table. Genes were retained if brain expression and maximum non-brain tissue expression were both nonzero and if brain expression was at least four-fold higher than the maximum expression observed in any other tissue. This yielded a set of 1,178 Ensembl gene IDs to form the brain-enriched set.

For each dataset-level object, gene detection was summarized from raw counts as total counts, mean counts per cell, number of cells with nonzero expression, and proportion of cells with nonzero expression from the complete gene universe for each dataset. A brain-enriched gene was considered detected at a given threshold if mean counts per cell was greater than zero and the proportion of cells with nonzero expression exceeded the threshold. Detection was evaluated at thresholds of *>* 0%, *>* 5%, *>* 10%, and *>* 30% of cells. Dataset-level values were summarized as the median number of detected brain-enriched genes across samples.

Gateway mOSN benchmarking used healthy donor samples only. FLEX v2 Gateway cells were subset to mature olfactory sensory neurons, and samples with more than 50 mOSNs were retained for detection analysis. External comparator objects were analyzed using the same gene-count summary and cell-count thresholds.

### Gene Ontology and UCell pathway scoring

Pathway activity was scored at the single-cell level using UCell, a rank-based method that computes gene-set enrichment within each cell (39). UCell scoring was applied to Gateway atlas cells using raw count matrices and the function `ScoreSignatures_UCell` with `maxRank = 2000`. Because UCell ranks genes within each cell, scores are less dependent on library size than average-expression module scores.

Gene sets were derived from MSigDB 2026.1 and included Gene Ontology Biological Process, Gene Ontology Cellular Component, Reactome, KEGG, Hallmark, olfactory, neuronal, compartmental, and control modules. For atlas-scale scoring, a neuronal function and compartment-focused gene-set panel was also constructed and scored with `ScoreSignatures_UCell` and `maxRank = 2000`. For external benchmarking, gene sets were rebuilt in Ensembl identifier space, and comparator datasets were scored using the same UCell framework.

For downstream summaries, UCell scores were aggregated by sample, cell type, dataset, and gene-set category using mean score, standard deviation, and the fraction of cells with positive scores. These summaries were z-scored to compare neuronal functional and compartmental programs across coarse-level cell types.

### Differential expression analysis

Cells were aggregated by summation to coarse cell-type and fine cell-type pseudobulks at the sample level, specifying donor and FLEX assay version for all FLEX v2 samples. A minimum of 200,000 UMIs per aggregate was used to define a valid pseudobulk per cell type.

Healthy controls were confirmed non-NDD via study record forms. Neurodegenerative donors were enrolled following clinical diagnosis of either Alzheimer’s disease or Parkinson’s disease by cognitive or motor-specialist neurologists. Three contrasts were evaluated: Alzheimer’s disease versus healthy control, Parkinson’s disease versus healthy control, and combined neurodegenerative disease versus healthy control. The combined neurodegenerative disease contrast grouped Alzheimer’s disease and Parkinson’s disease donors into a single disease class.

DESeq2 models used the design `~ age + sex + log_lib + group`, where `group` encoded disease status and healthy control was the reference level. `log_lib` was computed as log_10_ of the donor pseudobulk library size for the cell type of interest. Genes were not prefiltered before DESeq2. Independent filtering based on the `baseMean` cutoff was applied before Benjamini–Hochberg correction.

### Neurodegenerative disease gene visibility and dysregulation

We defined an a priori panel of commonly examined neurodegenerative disease-associated risk, therapeutic-target, and biomarker genes to assess baseline gene detectability and disease-associated dysregulation across Gateway olfactory epithelial cell types.

Baseline expression was determined in healthy controls. Disease-associated dysregulation of panel genes was assessed by differential expression analysis for Alzheimer’s disease, Parkinson’s disease, and combined neurodegenerative disease, defined as Alzheimer’s disease and Parkinson’s disease, contrasts.

Gene-set enrichment was performed from cell-type-specific DESeq2 Wald statistic results using fgsea (40). Enrichment was tested across an a priori defined panel of 33 pathways focused on neurodegeneration-relevant biology. fgsea was run with the non-default parameter `nPermSimple = 20000`.

## Data Availability

The single cell data for Gateway 4M are openly available on [CELLxGENE](https://cellxgene.cziscience.com/collections/8b35aa1f-6bcf-4a51-abc3-a3f336a44ae6). The available metadata for all fields is reflective of the reported values as completed by the participants. An experimental AI research assistant for interactive exploration of the expression data and analyses presented in this study is, at time of preprint release, available at chat.crownlands.com.

### Living and postmortem brain data acknowledgment

The results published here are in part based on data obtained from the AD Knowledge Portal (https://adknowledgeportal.org/). The Living Brain Project study participants are commended for their important role in science. The following centers, programs, departments and institutes within the Icahn School of Medicine at Mount Sinai supported the work: Department of Neurosurgery; Department of Genetics and Genomic Sciences; Department of Psychiatry; Department of Neuroscience; Department of Medicine; Friedman Brain Institute; Charles Bronfman Institute of Personalized Medicine; Hasso Plattner Institute for Digital Health; Black Family Stem Cell Institute; Pamela Sklar Division of Psychiatric Genomics; Mount Sinai Clinical Intelligence Center; Department of Psychiatry. The Living Brain Project is funded by the National Institute of Aging (R01AG069976) and the Michael J. Fox Foundation (Grant Number 18232). Analysis of the CommonMind Consortium data was performed with support from the National Institute of Mental Health (R01MH109897). Postmortem samples from Harvard Brain Tissue Resource Center and University of Miami Brain Endowment Brain were acquired under the National Institutes of Health NeuroBioBank request number 543. Postmortem samples from the New York Brain Bank of Columbia University were acquired under request number 1962. CommonMind Consortium data was utilized under the National Institutes of Mental Health Repository and Genomics Resource request identifier 5bffe32e97024. The following individuals are acknowledged for their advice and feedback over the course of the study: Elenita Sambat, Michael O. Hebb, Douglas M. Ruferfer, Wayne Goodman, Michael Donovan, Olha Fedoryshyn, Mary Fowkes, Vahram Haroutunian, Milind Mahajan, Sabina Berretta, Etty Cortes, Jean Paul Vonsattel, Eric J. Nestler, and Dennis S. Charney. This work is dedicated to Pamela Sklar.

## ACKNOWLEDGEMENTS

We thank Cindy Fang and Zach Gardell for their thoughtful comments and help constructing the article. This work could not have been possible without the physicians, clinic staff, and advisors who led and assisted in the studies. We extend sincere thanks to the people who participated in the clinical studies and donated samples to this research. We acknowledge the CZ CELLxGENE team for their assistance with the dataset and support of a key resource for the field.

